# Canonical and noncanonical TGF-β signaling regulate fibrous tissue differentiation in the axial skeleton

**DOI:** 10.1101/2020.11.04.368456

**Authors:** Sade W. Clayton, Ga I Ban, Cunren Liu, Rosa Serra

## Abstract

Previously, we showed that embryonic deletion of TGF-β type 2 receptor in mouse sclerotome resulted in defects in fibrous connective tissues in the spine. Here we investigated how TGF-β regulates expression of fibrous markers: Scleraxis, Fibromodulin and Adamtsl2. We showed that TGF-β stimulated expression of Scleraxis mRNA by two hours and Fibromodulin and Adamtsl2 mRNAs by eight hours of treatment. Regulation of Scleraxis by TGF-β did not require new protein synthesis; however, protein synthesis was required for expression of Fibromodulin and Adamtsl2 indicating the necessity of an intermediate. We subsequently showed Scleraxis was a potential intermediate for TGF-β-regulated expression of Fibromodulin and Adamtsl2. The canonical effector Smad3 was not necessary for TGF-β-mediated regulation of Scleraxis. Smad3 was necessary for regulation of Fibromodulin and Adamtsl2, but not sufficient to super-induce expression with TGF-β treatment. Next, the role of several noncanonical TGF-β pathways were tested. We found that ERK1/2 was activated by TGF-β and required to regulate expression of Scleraxis, Fibromodulin, and Adamtsl2. Based on these results, we propose a model in which TGF-β regulates Scleraxis via ERK1/2 and then Scleraxis and Smad3 cooperate to regulate Fibromodulin and Adamtsl2. These results define a novel signaling mechanism for TGFβ-mediated fibrous differentiation in sclerotome.

## INTRODUCTION

Fibrous tissues in the axial skeleton include the annulus fibrosus (AF) of the intervertebral disc (IVD), tendon, and ligaments. These fibrous tissues support movement of the axial skeleton, distribute mechanical load, and connect skeletal elements and muscle. Degeneration of AF is associated with disc disease and back pain. Loss of integrity of the AF can result in herniation of the IVD so that the nucleus pulposus (NP), the central component of the IVD, protrudes onto adjacent spinal nerves causing pain^1–3^. Fibrous connective tissues of the spine, including the AF, are not highly regenerative and damage results in significant pathology; nevertheless, how these tissues develop is not well-characterized even though this information would provide the basis of many therapeutic strategies.

The fibrous tissues of the spine are derived from sclerotome, an embryonic tissue composed of multipotent mesenchymal cells^4,5^ Previously, we showed that deletion of Tgfbr2 in Collagen2a (Col2a) expressing sclerotome led to significant defects in development of AF^6,7^. Others have shown that loss of Tgfbr2 in mesenchymal progenitors in mice results in defects in tendon formation, another fibrous tissue^8^. These studies indicate that TGF-β signaling is essential for development of fibrous tissues, but the mechanism of how TGF-β signaling promotes fibrous tissue differentiation in sclerotome remains to be elucidated.

TGF-β ligands transfer signals through type I and type II receptors (Tgfbr1 and Tgfbr2). Tgfbr2 binds the ligand and then recruits Tgfbr1 into a heterotetrameric complex. Tgfbr1 is activated upon phosphorylation by Tgfbr2 and then phosphorylates downstream effectors. The TGF-β receptor complex transduces signals through what are known as canonical and noncanonical pathways^9–12^. The canonical TGF-β signaling pathway uses Smad2 and/ or Smad3 to transfer signals. Smad2/3 are directly phosphorylated by Tgfbr1 and translocate to the nucleus to regulate gene transcription. One of the non-canonical TGF-β signaling pathways involves ERK/MAPK kinase. The ERK/MAPK pathway is induced by various stimuli. Activated ERK1/2 phosphorylates many proteins, including transcription factors, to regulate target genes and differentiation^13,14^. Previous studies showed that ERK signaling is associated with human skeletal diseases. For example, missense activating mutations in MEK1/2, a kinase upstream of ERK1/2, have been found in Cardio-facio-cutaneous syndrome^15^. In addition, haplo-insufficient expression of ERK2 was reported in DiGeorge syndrome^16^. These syndromes present with skeletal malformation, including short stature and limb abnormalities. Mice with loss of ERK1/2 in Col2a-positive cells (Col2a Cre-ERK1^-/-^; ERK2 ^f/f^) died immediately after birth, likely due to deformity of the rib cage. These mice also demonstrated kyphosis in the spine, which is also seen in mice with Tgfbr2 mutation^17^ suggesting ERK1/2 may have overlapping roles with Tgfbr2 in skeletal development.

Here, we address the mechanism of how TGF-β signaling affects fibrous tissue differentiation in sclerotome through canonical and non-canonical pathways. First, we showed that TGF-β regulates Scleraxis (Scx) mRNA, a transcription factor that regulates fibrous tissue differentiation, by 2hrs after treatment. In contrast, Fibromodulin (Fmod) and Adamtsl2 mRNAs, coding for ECM markers of fibrous tissue differentiation, were expressed later at 8hrs after treatment with TGF-β. Second, we showed that TGF-β-mediated expression of Scx mRNA did not require new protein synthesis. In addition, neither Smad2 nor Smad3 were required for TGF-β-mediated regulation of Scx. Next, we showed that new protein synthesis and Smad3 were required for TGF-β-mediated stimulation of Fmod and Adamtsl2 mRNAs. Nevertheless, exogenous Smad3 in conjunction with TGF-β treatment was not sufficient to super-regulate these markers suggesting another effector was involved. We then demonstrated that a non-canonical TGF-β signaling pathway involving ERK is required for expression of Scx, Fmod and Adamtsl2 mRNAs. Furthermore, we showed that Scx is required for full regulation of Fmod and Adamtsl2 in response to TGF-β. In summary, we have defined a new TGF-β signaling pathway in sclerotome that promotes differentiation of fibrous tissues in the axial skeleton.

## RESULTS

### TGF-β regulates fibrous tissue markers in the sclerotome

We previously identified, through microarray screens, several markers for fibrous tissues in the axial skeleton including Scx, Fmod, and Adamtsl2. These markers are enriched in the putative AF and regulated by TGF-β in sclerotome^18^. Here we further analyzed the timing of TGF-β-mediated induction of these gene’s mRNA after 2hrs and 8hrs of TGFβ1 treatment using quantitative real time RT-PCR (qPCR) analyzed using REST software. TGFβ1 induced Scx but not Fmod or Adamtsl2 after 2hrs of treatment. However, when cells were treated with TGFβ1 for 8hrs, expression of Fmod and Adamtsl2 was increased (Fig. 1A, Table S1). We then wanted to determine if TGF-β-mediated regulation of Scx, Fmod, and Adamtsl2 required new protein synthesis. Sclerotome cells were treated with cycloheximide (CHX) for 30 minutes to block protein synthesis, and then treated with vehicle control or TGFβ1 for 2hrs or 8hrs. A previous study using limb mesenchyme showed that Scx was induced after treatment with CHX, suggesting the existence of a labile protein repressor^19^. In addition, Scx mRNA was induced by TGF-β in the presence of CHX, suggesting direct regulation of Scx expression by TGF-β^19^. Similarly, in this experiment, Scx mRNA was increased in sclerotome in the presence of CHX and was induced after treatment with TGFβ1 in the presence of CHX (Fig. 1B, Table S2). The level of Scx mRNA induction after treatment with TGFβ1 in the presence of CHX was, however, attenuated relative to induction with TGFβ1 treatment alone (Fig. 1B, Table S2). Together the results indicate that TGF-β, at least in part, regulates Scx mRNA levels directly without new protein synthesis. In contrast, when cells were incubated with TGFβ1 for 8hrs in the presence of CHX, Fmod and Adamtsl2 were not regulated after TGFβ1 treatment (Fig. 1C, Table S3). The results indicate that new protein synthesis is required for TGF-β-meditated expression of Fmod and Adamtsl2 and suggest that an intermediate factor is involved in their regulation.

**Figure 1.**
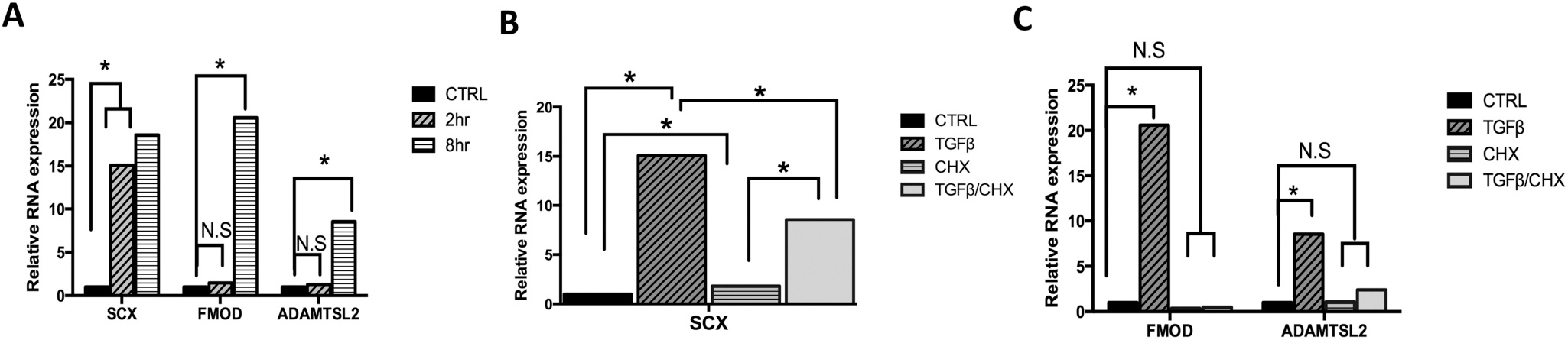
TGF-β regulates expression of fibrous tissue markers. (A) Primary sclerotome was treated with vehicle control or TGFβ1 for 2hrs or 8hrs. qPCR was used to determine the relative levels of Scx, Fmod and Adamtsl2 mRNA. (B, C) Sclerotome was treated with cycloheximide (CHX) for 30 mins then treated either with or without TGFβ1 for 2hrs (B) or 8hrs (C). Relative levels of fibrous tissue markers were determined by qPCR. All mRNA levels were normalized to Hprt and then compared to vehicle control using REST software. (*) indicates significance, p<0.05, n=3. Detailed results from qPCRREST analysis are shown in Table Sl-S3.

### Role of Smad-mediated signaling in regulation of fibrous tissue differentiation

Smad2 and Smad3 mediate TGF-β signaling by binding to regulatory regions of target DNA acting as transcription factors^10^. We tested the hypothesis that Smads are involved in TGF-β-mediated regulation of fibrous tissue differentiation by measuring expression of Scx, Fmod, and Adamtsl2 via qPCR while manipulating Smad2 and Smad3 activity. First, we determined the time course for activation of Smad2 and Smad3 after treatment with TGFβ1. Phosphorylated Smad2 and Smad3 (pSmad2 and pSmad3) were measured by western blot to show activation of signaling (Fig, 2A). Quantification shows that pSmad2 and pSmad3 were increased when sclerotome was treated with TGFβ1 after 2 and 8hrs while total levels of Smad2/3 were not altered (Fig 2B,C). We then used the Smad3 inhibitor, SIS3, and qPCR to determine if Smad3 was required for TGF-β-mediated regulation of the markers. SIS3 is known to specifically inhibit Smad3 and affects Smad3 target genes^20^. Primary sclerotome was pretreated with vehicle or two concentrations of SIS3 (5 and 10μM) for 24hrs at which time the cells were treated with TGFβ1 for 8hrs. Western blot confirmed that SIS3 inhibited activation of pSmad3 after treatment with 5 and 10μM TGFβ1 (Fig. 2D,E). Prg4 mRNA expression was also used as a positive control for SIS3 efficacy since we and others previously showed that Prg4 mRNA is regulated by Smad3^21^. Consistent with previous studies, inhibition of Smad3 with SIS3 blocked full induction of Prg4 by TGFβ1 in a dose dependent manner (Fig. 2F, Table S4). Induction of Fmod and Adamtsl2 by TGFβ1 were also attenuated following treatment of SIS3 (Fig. 2F, Table S4). These results suggest that Smad3 is required for TGF-β-mediated expression of Fmod and Adamtsl2. In contrast, SIS3 did not block TGF-β-mediated induction of Scx suggesting that regulation of Scx occurs through a non-canonical pathway or alternatively through pSmad2. Since it has been shown that Smad2 and Smad3 can regulate different downstream targets^22^, we tested a potential role for Smad2 in regulating Scx. Sclerotome was transduced with a dominant-negative form of Smad2 (Ad-DNSmad2). This point mutation in Smad2 was previously shown to dominantly attenuate signaling by both Smad2 and Smad3 due to its inability to be released from Tgfbr1^22^. Ad-DNSmad2 activity was confirmed as reduced pSmad2 and pSmad3 levels after treatment with TGF-β when compared to Ad-GFP control cells (Fig. 2G,H,I). As a positive control and as expected, infection with Ad-DNSmad2 attenuated induction of Adamtsl2 mRNA in response to TGF-β (Fig. 2J, Table S5). In contrast, Scx was still regulated by TGF-β to a statistically similar level as in the cells infected with the control (ad-GFP) virus (Fig. 2J, Table S5). The results suggested that TGFβ-mediated regulation of Scx did not require Smad2 or Smad3. Next, to determine whether exogenous Smad3 was sufficient to super-induce Scx, Fmod, or Adamtsl2 in the presence of TGFβ1, adenovirus encoding Smad3 (Ad-Smad3) was used to over-express Smad3 in sclerotome cultures. Immunoblot was used to confirm Smad3 overexpression in Ad-Smad3 infected cells (Fig. 2K, L). Cells were infected with Ad-Smad3 or Ad-GFP, as a control, for 48hrs and then treated with or without TGFβ1 for 8hrs. As expected, Prg4 mRNA levels were increased after treatment with TGFβ1 alone and super-induced in Ad-Smad3 infected cultures after Smad3 activation with TGFβ1 (Fig. 2M). In contrast, while TGFβ1 regulated expression of Scx, Fmod, and Adamtsl2, Smad3 did not super-induce expression after activation with TGFβ1 (Fig. 2M, Table S6). Since Smad3 was required for full TGF-β-mediated induction of Adamtsl2 and Fmod but not sufficient to super-induce expression after activation with TGFβ1, we propose an additional pathway is also required. This led us to investigate non-canonical signaling pathways.

**Figure 2.**
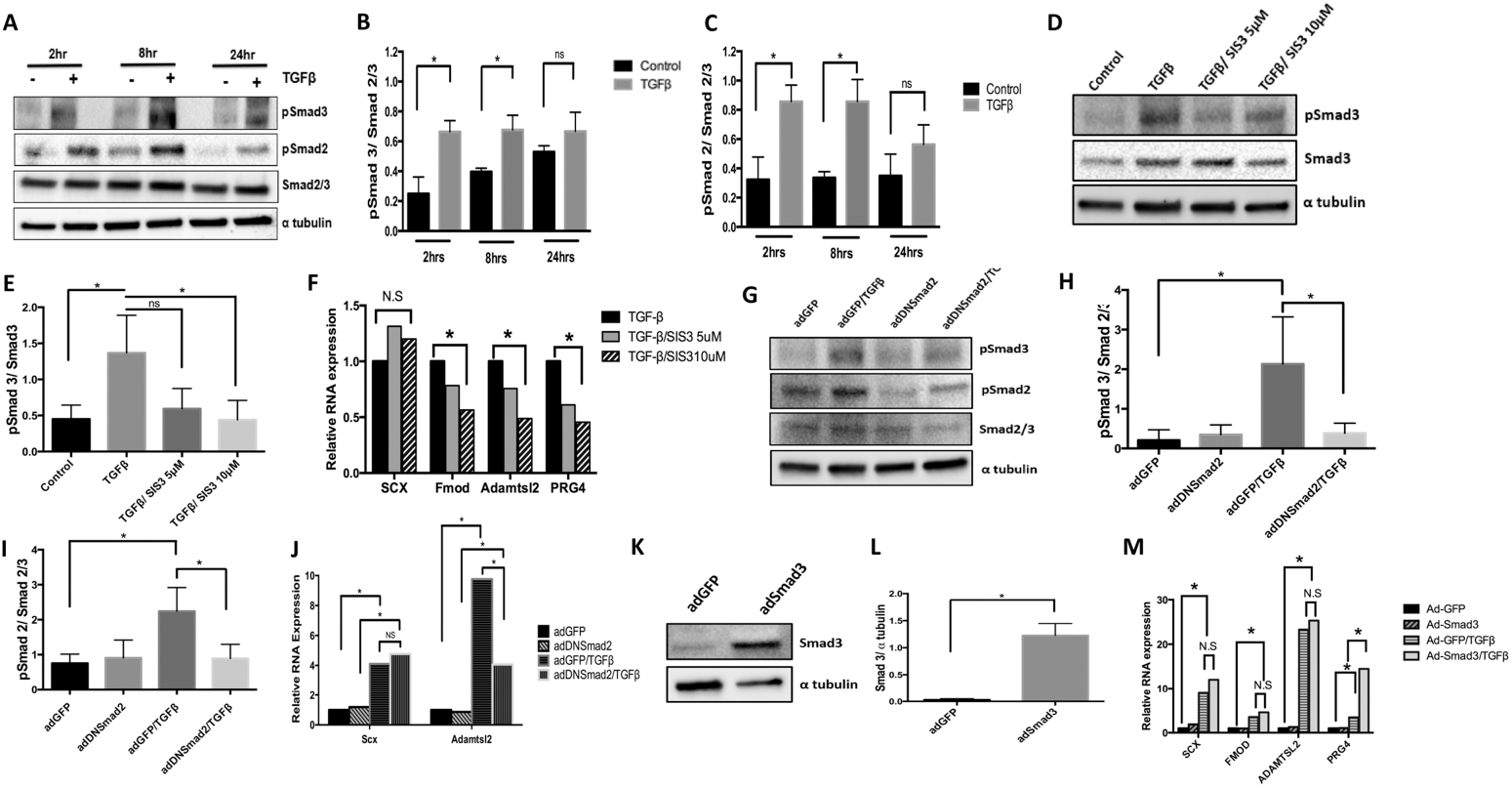
Smad3 is required for TGF-β to regulate expression of Fmod and Adamtsl2. (A) Primary sclerotome was treated with vehicle control or TGFβ1 for 2hrs, 8hrs or 24hrs. Immunoblot was used to determine the levels of pSmad3, pSmad2 and total Smad2/3. α-Tubulin was used as a general loading control. (B) Quantification of pSmad3 and (C) pSmad2 levels relative to total Smad2/3 are shown. Immunoblots were quantified using ImageJ. (*) indicates significance, p<0.05, n=3 for each (D) Sclerotome was pretreated with vehicle control or SIS3 (5μM and lOμM) for 24 hrs and then treated with or without TGFβ1 for 8hrs. pSmad3 and Smad2/3 protein levels were analyzed via immunoblot. α-Tubulin was used as a general loading control. (E) Smad3 activity was quantified from the immunoblots using ImageJ as the relative levels of pSmad3 normalized to total Smad2/3. (*) indicates significance, p<0.05, n=3 (F) Relative levels of Scx, Fmod, Adamtls2 and Prg4 mRNA were determined by qPCR after indicated treatment with SIS3. All mRNA levels were normalized to the housekeeping gene Hprt and then analyzed for significance using REST (*). Results are shown relative to TGFβ1 treated cells, n=3 (G) Cells were infected with AdDNSmad2 or AdGFP control virus 48 hours before cells were treated with or without TGFβ1 for 8 hrs. Immunoblot was used to visualize relative levels of pSmad3, pSmad2 and total Smad2/3. α-Tubulin is shown as a general loading control. (H) pSmad3 and (I) pSmad2 activity were quantified using ImageJ as the level of each phosphoSmad normalized to total Smad2/3. (*) indicates significance, p<0.05, n=3 (J) RNA was extracted from cells that were infected with AdDNSmad2 or AdGFP and treated or untreated with TGFβ1. Relative levels of Scx and Adamtsl2 mRNA were determined by qPCR. Expression was normalized to the housekeeping gene Hprt. Results are shown relative to the untreated control (*) indicates significance, p<0.05, n=5 (K) Cells were infected with AdSmad3 or AdGFP for 48 hrs and Smad3 expression was verified via immunoblot. (L) Immunoblots were quantified using ImageJ. (*) indicates significance, p<0.05, n=3 (M) RNA was isolated from cells infected with AdSmad3 or AdGFP for 48 hrs and then treated or untreated with TGFβ1 for 8hrs. Relative levels of Scx, Fmod, Adamtsl2, and Prg4 mRNA were determined by qPCR. Results are shown relative to Ad-GFP infected, untreated controls. All mRNA levels were normalized to the housekeeping gene Hprt. (*) indicates significance, p<0.05, n=3. Detailed results from qPCR REST analysis are shown in Table S4-S6. Immunoblots were cropped for clarity. Examples of uncropped blots are found in Supplementary Figures.

### Non-canonical pathways are regulated by TGF-β in the sclerotome

TGF-β transduces signals through not only Smad2/3 but also other kinases such as ERK1/2, p38, and AKT^12^. These Smad-independent pathways are often referred to as non-canonical TGF-β signaling pathways. To determine if TGF-β regulated fibrous tissue differentiation through non-canonical pathways, we determined the time course of activation of ERK1/2, p38, and AKT after treatment of sclerotome with TGFβ1 (Fig. 3A). Activated forms of ERK1/2 (pERK1/2) were observed after 2, 8, and 24hrs of TGFβ1 treatment (Fig. 3B). Total levels of ERK1/2 were not affected by TGFβ1. Activation of p38 as measured by the level of phosphorylated p38 (pp38) was observed at 2 and 8hrs of TGFβ1 treatment (Fig. 3C). Total p38 levels were not affected by TGFβ1 treatment. Increase in the phosphorylated form of AKT (p-AKT) was observed after TGFβ1 treatment for 2 and 8hrs (Fig. 3D). Total AKT was not affected. These data indicate that TGF-β can activate non-canonical signaling pathways in sclerotome.

**Figure 3.**
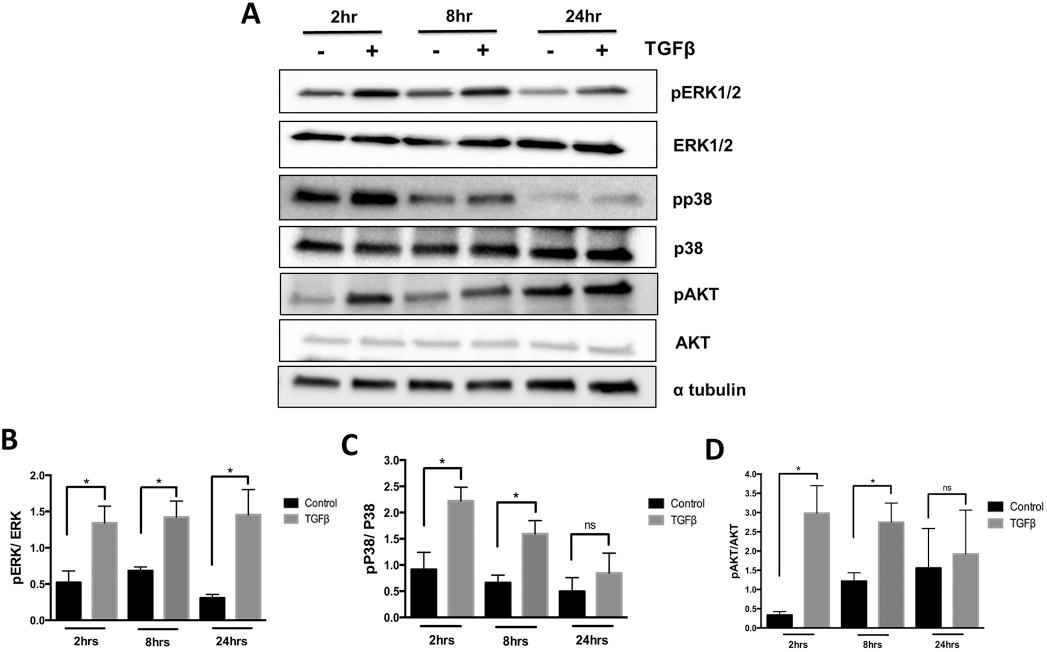
TGF-β signaling regulates noncanonical pathways in the sclerotome. Sclerotome was treated with vehicle control or TGFβ1 for 2, 8 or 24hrs. Immunoblot was used to determine activity of ERK, p38 and AKT. (B) ERK, (C) P38 and (D) AKT activity was quantified as the relative levels of the phosphoprotein over the total protein as determined from ImageJ scanned blots, α tubulin was used as a general loading control. (*) indicates significance, p<0.05, n=3. Immunoblots were cropped for clarity. Examples of uncropped blots are found in Supplementary Figures.

### p38 and AKT are not required for TGF-β-mediated regulation of fibrous tissue differentiation

We then tested whether activation of p38 is required for TGFβ-mediated fibrous tissue differentiation using a p38 antagonist, doramapimod (BIRB796), that inhibits all isoforms of p38^23^. TAK1/p38 signaling is activated by TGF-β in many cell types and mice with loss of the *Tak1* gene have a similar skeletal phenotype to mice deleted for *Tgfbr2*^24^. Isolated sclerotome was treated with BIRB796 or vehicle control, DMSO, for 24hrs and then treated with TGFβ1 or vehicle control for an additional 8hrs. The efficacy of the antagonist was measured by immunoblot of pMAPK-APK2, a direct downstream target of p38^25^. Activated MAPK-APK2 has multiple bands since p38 phosphorylates multiple residues to regulate activity^26,27^ (Fig. 4A). Treatment with TGFβ1 resulted in increased pMAPK-APK2 levels. Treatment with various concentrations of BIRB796 blocked the increase in pMAPK-APK2 in response to TGFβ1 (Fig. 4A,B). Using the same concentrations of BIRB796, we next measured Scx, Fmod, and Adamtsl2 expression in response to TGFβ1 using qPCR (Fig. 4C, Table S7). TGFβ1 was still able to up-regulate Scx, Fmod, and Adamtsl2 in the presence of the p38 antagonist suggesting that p38 signaling is not required for TGFβ1 to regulate fibrous tissue.

**Figure 4.**
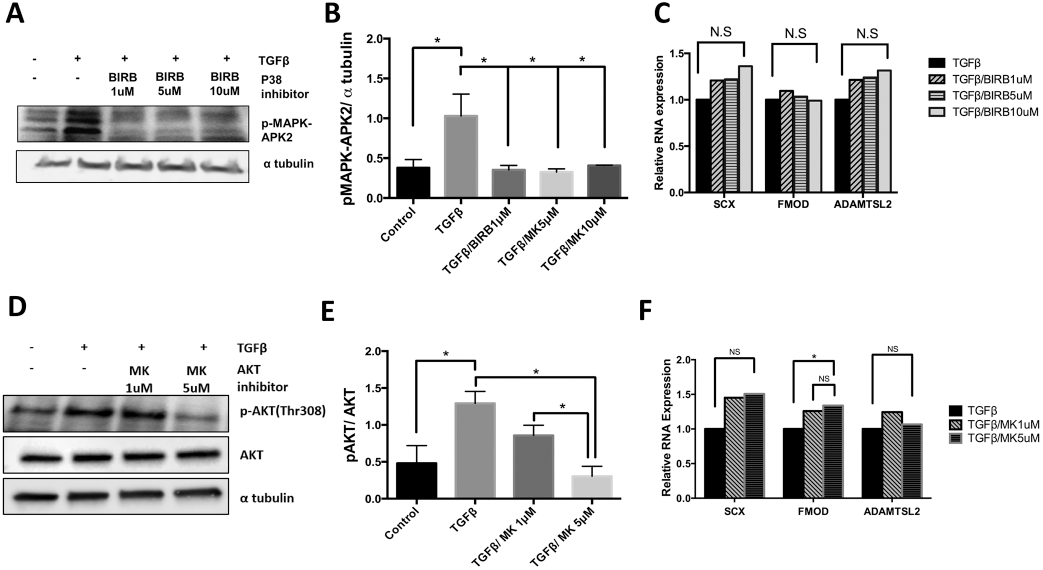
AKT and p38 are not required for TGF-β-mediated regulation of fibrous tissue markers. (A) Sclerotome was treated with varying concentrations (lμM, 5μM, 10μM) of P38 inhibitor, BIRB, for 24hrs and then treated with TGFβ1 for 8hrs. Immunoblot was used to determine relative levels of the p38 target, p-MAPK-APK2. α-tubulin was used as a loading control. (B) p-MAPK-APK2 levels were quantified using Image J and GraphPad Prism. (*) indicates significance, p<0.05, n=3. (C) RNA was isolated from cells treated with BIRB and TGFβ1 as indicated. qPCR was used to measure the relative expression of Scx, Fmod, and Adamtsl2. mRNA levels were normalized to the housekeeping gene HPRT. Expression is shown relative to the TGFβ1 treated control. Results were analyzed with REST. (*) indicates significance, p<0.05, n=3. (D) Sclerotome was treated with varying concentrations (lμM, 5μM) of AKT inhibitor, MK, for 24hrs and then treated with TGFβ1 for 8hrs. Immunoblot was used to measure relative levels of pAKT and total AKT. α-tubulin was used as a general control. (E) Immunoblots were scanned using Image J and quantified. Activation of AKT was measured as pAKT over total AKT. (*) indicates significance, p<0.05, n=3 (F)mRNA was isolated from cells treated with MK and TGFβ1 as indicated. qPCR was used to measure the relative levels of Scx, Fmod, or Adamtsl2 mRNA. All mRNA levels were normalized to the housekeeping gene HPRT. Expression is shown relative to the TGFβ1 treated samples. Results were analyzed with REST. (*) indicates significance, p<0.05,n=3. Detailed results from qPCR REST analysis are shown in Table S7, S8. Immunoblots were cropped for clarity. Examples of uncropped blots are found in Supplementary Figures.

We then tested whether activation of AKT signaling was required for TGF-β-mediated fibrous tissue differentiation by using an AKT inhibitor, MK2206^28^. The PI3K/AKT pathway is known to have an important role in cartilage development and is also activated by TGF-β in other cell types^29^. The concentration of MK2206 necessary to block activity was determined by immunoblots of the active form of AKT. As expected, MK2206 attenuated TGF-β-induced phosphorylated AKT in the sclerotome (Fig. 4D,E). Sclerotome was then treated with MK2206 and TGFβ1 so that the effects of AKT on fibrous tissue differentiation could be determined as expression of fibrous tissue markers by qPCR. mRNA levels of fibrous tissue markers were the same after treatment with TGF-β in the presence or absence of the AKT inhibitor, MK2206 (Fig. 4F, Table S8). These results suggest that AKT was not involved in TGF-β-mediated induction of fibrous tissue differentiation.

### Role of ERK1/2 in fibrous tissue differentiation and TGFβ-mediated inhibition of chondrogenesis

Next, we investigated whether ERK1/2 signaling mediates the effect of TGFβ1 on fibrous tissue differentiation. We used a MEK inhibitor, PD184352, to inhibit ERK1/2 activity^30^. MEK is directly upstream of ERK1/2 and inhibition of MEK is a common method to inhibit ERK1/2 activity. Mice with deletion of ERK1 and ERK2 in their axial skeletons demonstrate kyphotic deformity in the thoracic spine^17^. To determine the necessary drug concentration, sclerotome was treated with PD184352 in presence or absence of TGFβ1. Treatment with TGFβ1 resulted in an increase in phosphorylated ERK1/2, as previously shown (Fig. 5A,B). Efficacy of 1, 5 and 10μg/ml doses of PD184352 was confirmed via immunoblot as a dose dependent reduction in the levels of TGFβ1-induced phosphorylated ERK1/2 (pERK1/2) (Fig. 5A,B). Next, expression of markers for fibrous tissue differentiation were measured via qPCR after TGFβ1 treatment. Induction of Scx, Fmod, and Adamtls2 by TGF-β were all attenuated in a dose dependent manner after treatment with PD184352 (Fig. 5C, Table S9). The results indicated that ERK1/2, but not p38 or AKT, is required for TGFβ1-mediated fibrous tissue differentiation. Next, we tested the specificity of the response and determined whether TGFβ1 utilizes ERK1/2 signaling to down-regulate the expression of early markers for chondrogenesis, the alternate cell fate of sclerotome. Previously, our laboratory showed that TGFβ1 inhibits chondrogenesis and down-regulates EBF1 and cMAF, vertebrae enriched genes that are involved in chondrogenesis, in the sclerotome within 8hrs of treatment^31^. EBF1 and cMAF mRNA levels were measured by qPCR after treatment with TGFβ1 and PD184352. Inhibition of ERK1/2 did not affect TGFβ1-mediated down-regulation of these genes, indicating that ERK1/2 signaling is specific for TGFβ-mediated regulation of fibrous differentiation (Fig. 5D, Table S10).

**Figure 5.**
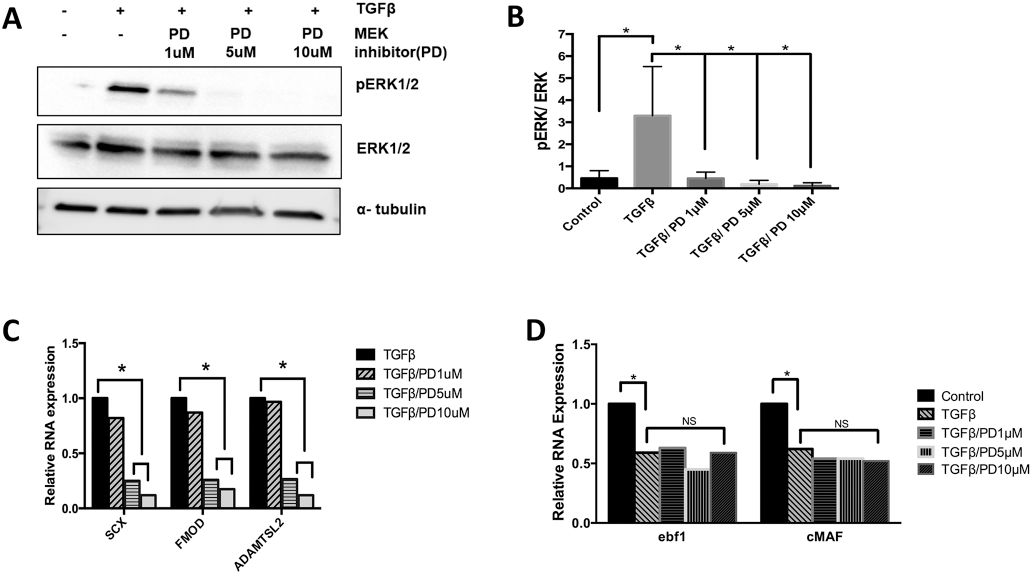
ERK is required for fibrous tissue marker regulation but not required to inhibit chondrogenesis. (A) Sclerotome was treated with a MEK inhibitor PD184352, PD,to inhibit ERK activity, for 24hrs and then cells were treated with TGFβ1 for 8hrs. Immunoblot was used to determine relative levels ofpERKl/2, ERK1/2, and α-tubulin. (B) Blots were quantified using ImageJ and Graphpad Prism. Activity of ERK was determined as the ratio of pERK over ERK under the indicated conditions. (*) indicates significance, p<0.05,n=3. (C, D) RNA was isolated from cells under the indicated conditions and qPCR was used to determine the relative expression of (C) Scx, Fmod and Adamtsl2 mRNA or (D) Ebfl and cMAF mRNA. mRNA levels were normalized to Hprt. Expression is shown relative to the untreated control. Data analyzed by REST software. (*) indicates significance, p<0.05,n=3. Detailed results from qPCR REST analysis are shown in Table S9, S10. Immunoblots were cropped for clarity. Examples of uncropped blots are found in Supplementary Figures.

### Scx is required for TGFβ-mediated regulation of Fmod and Adamtsl2

Finally, we tested the hypothesis that Scx is involved in subsequent TGFβ-mediated expression of the ECM fibrous markers Fmod and Adamtsl2. Scx is a basic helix loop helix (bHLH) transcription factor^8^ that is regulated earlier than the other fibrous tissue markers (Fig. 2A) and is up-regulated at least in part by TGF-β even in the absence of new protein synthesis suggesting direct regulation by TGF-β (Fig. 2B). To determine if Scx expression was required to regulate Fmod and Adamtsl2, we inhibited Scx synthesis by transducing C_3_H_10_T_1/2_ cells, a murine mesenchymal stem cell line, with 30pM of Scx specific siRNA or a 30pM of Scrambled (Scram) siRNA as a control. We and others have previously shown that fibrous genes are regulated by TGFβ1 in C_3_H_10_T_1/2_ cells^32,33^. Western blot analysis showed a significant decrease in Scx protein expression in Scx siRNA transduced cells when compared to Scram control (Fig. 6A,B). qPCR determined that mRNA levels of Scx were also significantly decreased with Scx siRNA treatment (Fig. 6C, Table S11). We then wanted to test whether a decrease in Scx expression affected expression of Fmod and Adamtsl2. As expected, qPCR analysis determined that in Scram control siRNA transduced cells treatment with TGFβ1 resulted in an increase in marker expression. Basal Fmod expression was significantly downregulated in Scx siRNA transduced cells when compared to Scram siRNA suggesting Scx is important for regulation of Fmod in general. Furthermore, Fmod was not up-regulated by TGFβ1 in Scx siRNA transduced cells, signifying that Scx expression is also required for TGF-β mediated regulation of Fmod (Fig. 6D, Table S12). Inhibtion of Scx with siRNA did not affect basal levels of Adamtsl2 mRNA, but, induction of Adamtl2 by TGFβ1 was blocked, indicating that Scx is also required for TGF-β mediated regulation of Adamtsl2 (Fig. 6D, Table S12).

**Figure 6.**
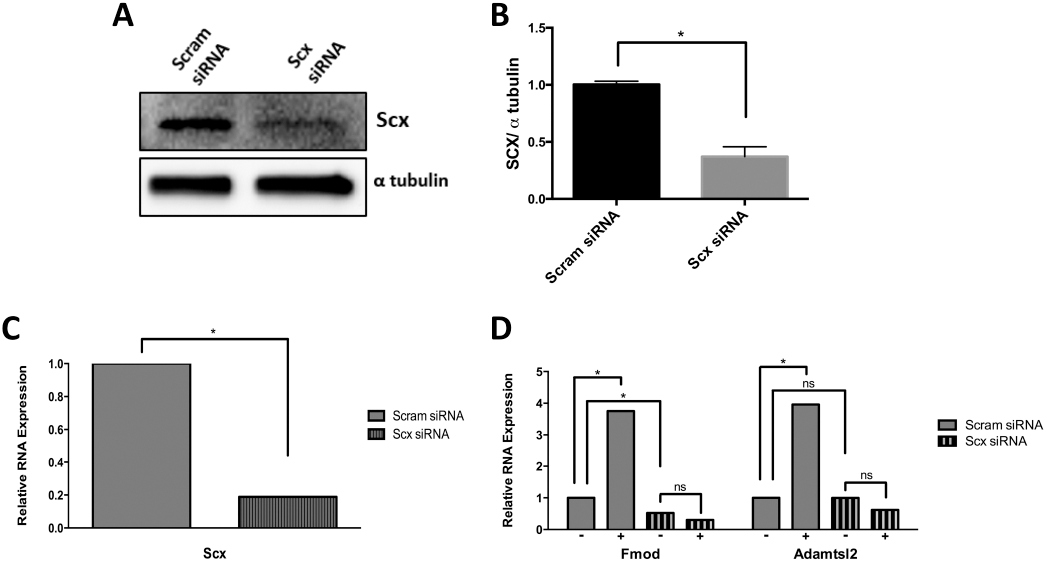
Scx is required for the expression of Fmod and Adamtsl2. (A) C_3_H_10_T_1/2_ cells were transduced with a scrambled siRNA control (Scram) or Scx specific siRNA for 24hrs. Representative immunoblot showing Scx protein, α-tubulin is used as a loading control. (B) Quantification of Scx immunoblots comparing Scram transduced and Scx siRNA transduced cells. Protein levels were normalized to α tubulin and quantified using ImageJ and GraphPad Prism, n=3, (*) indicates significance, p<0.05. (C) Scx mRNA levels were determined by qPCR in Scram transduced and Scx siRNA transduced cells. Data was normalized to HprtmRNA and analyzed using REST software, n=6, (*) indicates significance, p<0.05. (D) RNA was collected from cells that were transduced with Scram siRNA or Scx siRNA for 24 hrs and subsequently either left untreated (-) or treated with TGFβl(+) for 8hrs. Fmod and Adamtsl2 mRNA levels were determined by qPCR. mRNA levels were normalized to Hprt. qPCR data was analyzed by REST software, n=6, (*) indicates significance, p<0.05. Detailed results from qPCR REST analysis are shown in Table SI 1, S12. Immunoblots were cropped for clarity. Examples of uncropped blots are found in Supplementary Figures.

## DISCUSSION

We previously identified several TGF-β regulated genes for which expression is enriched in the putative IVD region of the embryonic sclerotome, including Adamtsl2, Fmod, and Scx^18,31,34^. Localization to a patterned subset of sclerotome was previously confirmed by *in-situ* hybridization^18,35^. Mutations in Adamtls2 result in a condition called Geleophysic dysplasia, which is characterized in part by short stature and joint contractures. It was recently shown that the joint contractures and short limb phenotype were mediated by Adamtsl2 expression in Scx-positive cells^35^. Phenotype in the spine was not discussed. Likewise, Fmod is required for development of tendons and other fibrous tissues^36^. Almost nothing is known about upstream regulators of Adamtsl2 or Fmod expression. We previously showed that both are strongly regulated by TGF-β, which is also involved in development of tendon and other fibrous tissues^8,18,31,34^. Here we show that Fmod and Adamtsl2 mRNAs are regulated by non-canonical and canonical signaling pathways, ERK1/2 and Smad3. Furthermore, experiments using a protein synthesis inhibitor, CHX, showed new protein synthesis was required for TGF-β-mediated upregulation of Fmod and Adamtsl2 mRNAs. These results suggest that an intermediary protein must be made so that TGF-β can regulate the expression of these ECM proteins. Scx, a basic helix loop helix (bHLH) transcription factor^8^, was up-regulated, at least in part, by TGF-β even in the absence of new protein synthesis suggesting direct regulation by TGF-β. Induction of Scx by TGF-β in CHX treated cells was, however, attenuated suggesting that there may be two mechanisms for TGF-β-mediated regulation of Scx. One that does not require de novo protein synthesis, perhaps regulating the immediate response to TGF-β, and another that requires new protein, perhaps regulating maintenance of Scx expression in fibrous cells. In addition, Scx required ERK1/2 but not Smad3 for TGF-β-mediated regulation. We then showed that Scx was required to regulate Fmod and Adamtsl2. We propose that TGF-β regulates Fmod and Adamtsl2 through induction of one or more transcription factors, including Scx.

Deletion of Scx results in alterations in the development of long tendons of the spine^37^. More recently, it was shown that loss of Scx disrupts maturation of AF, and the loss of Sox9 in Scx expressing cells disrupts formation of the AF and chondro-tendinous/ligamentous junctions^38,39^. Scx expression is up-regulated by TGF-β in mesenchymal cells including primary sclerotome and C_3_H_10_T_1/2_ cells and down-regulated in tissues in which Tgfbr2 has been conditionally deleted^8,31,34^. bHLH transcription factors like Scx generally bind to DNA as dimers to elements known as E-boxes^40^. It has also been shown that Scx can bind to Smad3 to regulate down-stream gene expression^41,42^.

A little is known about how Scx expression is regulated. FGF was the first factor shown to regulate Scx during development^43^. It was shown that FGF secreted from the myotome acted on the adjacent sclerotome to induce Scx expression and fibrous differentiation. FGF is known to signal though ERK1/2 and it was shown that ERK1/2 is required for Scx expression in chick axial and appendicular skeleton^43–45^. TGF-β is known to activate ERK1/2 through a non-canonical pathway^46,47^. In this study we showed that ERK1/2 is required for TGF-β-mediated expression of Scx; however canonical signaling through Smad was not necessary to regulate Scx mRNA in sclerotome. Likewise, regulation of Scx in cardiac fibroblasts is Smad3-independent^48^ and Scx expression is not depleted in tendons of embryonic *Smad3* knock out mice^41^. However, in contrast to these observations, one study showed that TGF-β-mediated expression of Scx in limb was independent of ERK1/2 and dependent on Smad3^45^. This discrepancy may be due to different cell type, microenvironment, or developmental stage.

Since Smad3 was necessary but not sufficient to super-induce expression of Fmod and Adamtsl2, we hypothesized that TGF-β likely uses a combination of signaling pathways in addition to Smad3 (Fig. 7). Noncanonical TGF-β signaling pathways, which are Smad-independent, use various kinases like p38, AKT and ERK1/2^12^. TGF-β differentially activates these non-canonical pathways based on cell type and other circumstances. Previous studies showed that p38 mediates TGF-β signaling in articular chondrocytes^49^. Also, mice lacking TAK1, which lies upstream of p38, have defects in joint formation, similar to that observed in the absence of Tgfbr2^24^. However, the results here indicated that TGF-β does not utilize p38 signals to regulate fibrous tissue differentiation. Next, we tested the role of ERK1/2, since ERK1/2 is also known to mediate TGF-β signaling^12^. Results indicated that ERK1/2 plays an important role in mediating fibrous tissue differentiation by TGF-β. Inhibition of ERK1/2 did not alter the response of cartilage genes to TGF-β treatment, indicating that the role of ERK1/2 is specific to fibrous tissue differentiation.

**Figure 7.**
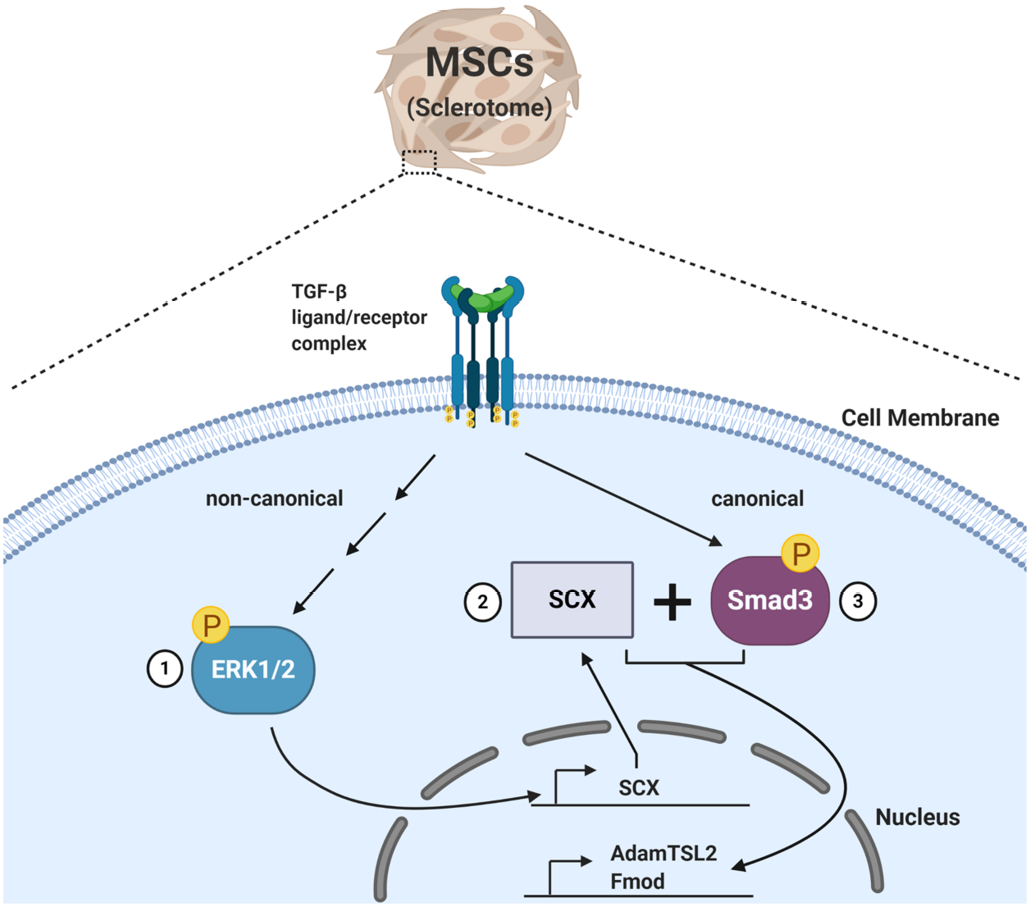
A schematic of the proposed mechanism of TGF-β-mediated regulation of fibrous tissue differentiation from sclerotome. We hypothesize that TGFβ regulates fibrous tissue differentiation by first regulating non-canonical signaling through ERK1/2 (1), which is required for Sex mRNA expression (2). We propose that subsequently, Sex and Smad3 cooperate to regulate the ECM markers, Fmod and Adamtsl2 (3).

This study demonstrates that canonical and noncanonical TGF-β signaling pathways regulate fibrous tissue differentiation in sclerotome. We propose that TGF-β acting through ERK1/2 regulates expression of Scx. Subsequently, Scx and Smad3 cooperate to regulates expression of Fmod and Adamtsl2 (Fig 7). This study begins to define a novel signaling pathway for TGF-β in sclerotome that regulates fibrous tissue differentiation.

## MATERIALS AND METHODS

All experiments with animals were approved by the University of Alabama Institutional Animal Care and Use Committee (IACUC) under protocol number 09713. All experiments with animals and all methods were carried out in accordance with relevant guidelines and regulations.

### Sclerotome isolation and cell culture

Sclerotome was isolated from wild type mouse embryos at E11.5 days as previously described^18^. Embryos were generated by mating C57BL/6 male mice to ICR female mice. The tissue was digested with Collagenase D, dissociated into individual cells, and plated. Cells were plated in high cell density micromass cultures with 3.0 X 10^5^ cells per 20ul drop of culture medium (40% DMEM, 60% Ham’s F-12, 10% heat-inactivated FBS, 0.5mM L-glutamine, 1% Penicillin/Streptomycin, 0.25mM Ascorbic Acid and 1mM β-glycerophosphate). Five to six micromasses were plated onto one 35mm culture dish (Corning). Cells in micromass drops were treated with varying dosages of inhibitors as indicated, SIS3 (EMD Millipore), PD184352 (Sigma Aldrich), BIRB 796 (Selleck Chemicals) or MK 2206 (Selleck Chemicals) and plated for one hour to allow adherence. After one hour, culture dishes were flooded with medium that contained the indicated inhibitors. Micromass cultures were then maintained at 37°C in a humidified incubator with 5% CO2 for 24hrs with the inhibitors before treatment with 5ng TGFβ1/ml (R&D Systems) for the indicated times.

### Adenoviruses

Adenoviruses were used to both overexpress Smad3 and inhibit Smad2/3 (via dominant-negative Smad2). To overexpress Smad3 in sclerotome cells, an adenovirus encoding a FLAG-tagged Smad3 protein with an enhanced green fluorescent protein (eGFP) reporter (Ad-Smad3) or control virus (Ad-GFP) was used. To inhibit Smad2/3 regulation, a dominant-negative Smad2 virus (Ad-DNSmad2) that blocks both Smad2 and Smad3 signaling was used. The generation of the Ad-DNSmad2 virus was described previously^22,50,51^. Both the Ad-Smad3 and Ad-DNSmad2 viruses were gifts from Dr. Rik Derynck^22^. Adenoviruses were amplified in and collected from

HEK293T cells. Viral titers were determined by counting the number of eGFP positive cells (for Ad-GFP, Ad-Smad3) or using immunofluorescence to count the number of Hexon, 1:1000 Catalog #0400-0064, (Bio-Rad Laboratories) positive cells labeled with Alexa488, 1:1000 Catalog #A11006, (Thermo Fisher Scientific) secondary antibody (for Ad-DNSmad2). Cells were mixed with 50 M.O.I (Multiplicity of Infection) of virus, plated in micromass cultures for 1hr and then incubated for 48hrs with the various adenoviruses before treatment with 5ng TGFβ1/ mL (R&D Systems) for the indicated times.

### Quantitative real-time RT-PCR (qPCR)

Cultures were harvested using TRIzol reagent (Thermo Fisher Scientific) and total RNA extracted using a Direct-zol RNA MiniPrep kit (Zymo Research). The concentration of RNA was measured with a spectrophotometer (Nano Drop) and 10ng of RNA per sample was used in each reaction according to instruction in QuantiFast SYBR Green RT-PCR kit (QIAGEN). Samples were subsequently loaded on to a LightCycler 480 Instrument II (Roche). Crossing point (Cp) values were obtained and converted to relative mRNA expression levels using the Relative Expression Software Tool (REST) 2009 (Qiagen)^52^. REST analysis provides statistical testing for nonparametric data using a pair-wise fixed reallocation randomization test to determine significance (p > 0.05). “n” denotes the number of biological replicates. HPRT was used as a reference/normalization control. Primer sequences for qPCR are shown in Table S13.

### Western blot

Cultured sclerotome was lysed with RadioImmunoPrecipitationAssay (RIPA) buffer containing phosphatase and protease inhibitors (Roche). The concentration of total protein was measured using a DC Protein Assay kit (Bio-Rad Laboratories). 25-30μg of protein lysate per sample was loaded in 4-20% polyacrylamide gels (Bio-Rad Laboratories) and separated protein was transferred from gels to polyvinylidene fluoride membranes (PVDF) using a Trans-Blot Turbo Transfer system (Bio-Rad Laboratories). The membrane was blocked with 5% Bovine Serum Albumin (Sigma-Aldrich) and incubated with various primary antibodies. Membranes were then washed with Tris-buffered saline containing 0.1% Tween 20 (TBST) and incubated with various secondary antibodies. The chemiluminescent detection of the horseradish peroxidase on the secondary antibodies was detected by the Supersignal West Dura kit (Thermo Scientific). Images and of western blots were acquired on a ChemiDoc MP system (Bio-Rad Laboratories) and quantification of blots was performed on ImageJ. All statistical analyses of immunoblots was performed using GraphPad Prism either using an unpaired t-test or a one-way ANOVA with a Tukey’s post hoc analysis. All immunoblots are n=3 (biological replicates) and asterisks denote p < 0.05.

### Antibodies

For western blot analysis, the following primary antibodies were used at a dilution of 1:1000 unless stated otherwise: pSmad2 Catalog #3104S (Cell Signaling), pSmad3 Catalog #9520S (Cell Signaling), Smad2/3 Catalog #3102S (Cell Signaling), Smad3 Catalog #9513S (Cell Signaling), pERK1/2 Catalog #9101S (Cell Signaling), ERK1/2 Catalog #4695S (Cell Signaling), pp38 Catalog #9211S (Cell Signaling), p38 Catalog #9212S (Cell Signaling), pAKT Catalog #13038S (Cell Signaling), AKT Catalog #9272S (Cell Signaling), pMAPKAPK2 Catalog #3007S (Cell Signaling), SCXA Catalog #PA5-23943 (Invitrogen), and Alpha tubulin Catalog #200-301-880, 1:5000 (Rockland). The following secondary antibodies were used at a dilution of 1:1000: anti-Rabbit-HRP, Catalog #7074S (Santa Cruz Biotechnology) and antimouse Dylight 649, Catalog #610-143-003 (Rockland).

### siRNA

Murine C_3_H_10_T_1/2_ cells (ATCC) were plated in 6-well tissue culture treated plates (Corning Costar) at a density of 3×10^5^ cells/well in cell culture medium (88% DMEM, 10% heat-inactivated FBS, 1% 0.5mM L-glutamine, 1% Penicillin/Streptomycin,) for 24hrs. After 24hrs, culture medium was replaced with MEM Alpha 1X (Gibco) and cells were transfected using Lipofecatime RNAiMax reagent (ThermoFisher Scientific) with either 30pM of Scleraxis siRNA Catalog #sc-153257 (Santa Cruz) or 30pM of Scrambled Control siRNA (FITC Conjugate)-A Catalog #sc-36869 (Santa Cruz) for 24hrs. Cells were then lysed with RIPA Buffer for western blot analysis or TRIzol reagent for qPCR. Western blot data n=3 and qPCR data n=6. “n” denotes number of biological replicates.

## Acknowledgements

This study was funded by R01 AR053860 to R.S and T32 AR069516 (PI Bridges) to SWC.

## Author Contributions

S.W.C., G.I.B., and R.S. contributed to the conception and design of the study. S.W.C., G.I.B., and C.L. acquired the data. S.W.C., G.I.B., and R.S. contributed to the analysis and interpretation of the data. S.W.C., G.I.B. and R.S. wrote the manuscript. All authors approved the final version of the manuscript and take responsibility for the integrity of the work.

