## Supplemental for "Canonical and noncanonical TGF-β signaling regulate fibrous tissue differentiation in the axial skeleton"

Table S1. qPCR details for Figure1A. Data was normalized to HPRT and expression levels calculated relative to vehicle control.

| Comparison | Fold Change | Confidence Intervals 65% | P-value | Result |
|---|---|---|---|---|
| SCX | | | | |
| Control (Vehicle) | 1 | 1 | | |
| Control vs T_2hr | 15.086 | 12.742 - 18.183 | 0.0001 | UP |
| Control vs T_8hr | 18.586 | 15.587 - 22.730 | 0.0001 | UP |
| FMOD | | | | |
| Control (Vehicle) | 1 | 1 | | |
| Control vs T_2hr | 1.462 | 0.282 - 11.520 | 0.609 | - |
| Control vs T_8hr | 20.604 | 15.005 - 28.415 | 0.0001 | UP |
| ADAMTSL2 | | | | |
| Control (Vehicle) | 1 | 1 | | |
| Control vs T_2hr | 1.282 | 0.220 - 10.591 | 0.782 | - |
| Control vs T_8hr | 8.560 | 3.884 - 18.177 | 0.0001 | UP |

Table S2. qPCR details for Figure1B. Data was normalized to HPRT and expression levels calculated relative to vehicle control.

| Comparison | Fold Change | Confidence Intervals | P-value | Result |
|---|---|---|---|---|
| | | 65% | | |
| SCX | | | | |
| Control (Vehicle) | 1 | 1 | | |
| Control vs T | 15.086 | 12.742 - 18.183 | 0.0001 | UP |
| Control vs CHX | 1.820 | 1.430 - 2.236 | 0.0001 | UP |
| Control vs T/CHX | 8.554 | 5.799 - 10.936 | 0.0001 | UP |
| CHX vs T/CHX | 4.700 | 3.192 - 6.453 | 0.0001 | UP |
| T vs T/CHX | 0.567 | 0.396 – 0.734 | 0.0001 | DOWN |

Table S3. qPCR details for Figure1C. Data was normalized to HPRT and expression levels calculated relative to vehicle control.

| Comparison | Fold Change | Confidence Intervals | P-value | Result |
|---|---|---|---|---|
| | | 65% | | |
| FMOD | | | | |
| Control (Vehicle) | 1 | 1 | | |
| Control vs T | 20.604 | 15.005 - 28.415 | 0.0001 | UP |
| Control vs CHX | 0.356 | 0.249 - 0.487 | 0.0001 | DOWN |
| Control vs T/CHX | 0.495 | 0.374 - 0.636 | 0.0001 | DOWN |
| CHX vs T/CHX | 1.388 | 1.028 - 1.928 | 0.016 | UP |
| ADAMTSL2 | | | | |
| Control (Vehicle) | 1 | 1 | | |
| Control vs T | 8.560 | 3.884 - 18.177 | 0.0001 | UP |
| Control vs CHX | 1.102 | 0.481 - 1.972 | 0.694 | - |
| Control vs T/CHX | 2.402 | 1.101 - 3.907 | 0.005 | UP |
| CHX vs T/CHX | 2.180 | 1.521 - 3.301 | 0.0001 | UP |

Table S4 qPCR details for Figure 2F. SCX, FMOD, ADAMTSL2 or PRG4 data was normalized to HPRT, expression levels calculated relative to TGFβ treated.

| Comparison | Fold Change | Confidence Intervals | P-value | Result |
|---|---|---|---|---|
| | | 65% | | |
| SCX | | | | |
| TGFβ | 1 | 1 | | |
| T vs T/SIS3 5ug/ml | 1.314 | 0.056 - 29.668 | 0.624 | - |
| T vs T/SIS3 10ug/ml | 1.201 | 0.064 - 21.577 | 0.709 | - |
| FMOD | | | | |
| TGFβ | 1 | 1 | | |
| T vs T/SIS3 5ug/ml | 0.783 | 0.679 - 0.901 | 0.034 | DOWN |
| T vs T/SIS3 10ug/ml | 0.565 | 0.440 - 0.754 | 0.0001 | DOWN |
| ADAMTSL2 | | | | |
| TGFβ | 1 | 1 | | |
| T vs T/SIS3 5ug/ml | 0.757 | 0.666 - 0.861 | 0.000 | DOWN |
| T vs T/SIS3 10ug/ml | 0.488 | 0.368 - 0.647 | 0.003 | DOWN |
| PRG4 | | | | |
| TGFβ | 1 | 1 | | |
| T vs T/SIS3 5ug/ml | 0.610 | 0.409 - 1.017 | 0.144 | - |
| T vs T/SIS3 10ug/ml | 0.455 | 0.242 - 0.860 | 0.048 | DOWN |

Table S5. qPCR details for Figure 2J. SCX or ADAMTSL2 data was normalized to HPRT, expression levels calculated relative to vehicle control.

| Comparison | Fold Change | Confidence Intervals 65% | P-value | Result |
|---|---|---|---|---|
| SCX | | | | |
| Control (Vehicle) | 1 | 1 | | |
| Ad-GFP vs Ad-DNSmad2 | 1.192 | 0.824 - 1.740 | 0.366 | - |
| Ad-GFP vs Ad-GFP/T | 4.105 | 3.147 - 5.463 | 0.002 | UP |
| Ad- DNSmad2 vs Ad-DNSmad2/T | 4.769 | 2.667 - 6.743 | 0.004 | UP |
| Ad-GFP/T vs Ad-DNSmad2/T | 1.162 | 0.769 - 2.287 | 0.743 | - |
| ADAMTSL2 | | | | |
| Control (Vehicle) | 1 | 1 | | |
| Ad-GFP vs Ad-DNSmad2 | 0.868 | 0.622 - 1.240 | 0.449 | - |
| Ad-GFP vs Ad-GFP/T | 9.758 | 5.950 - 17.070 | 0.000 | UP |
| Ad- DNSmad2 vs Ad-DNSmad2/T | 4.066 | 2.219 - 8.230 | 0.010 | UP |
| Ad-GFP/T vs Ad-DNSmad2/T | 0.417 | 0.182 - 0.846 | 0.040 | DOWN |

Table S6. qPCR details for Figure 2M. SCX, FMOD, ADAMTSL2 and PRG4 data was normalized to HPRT, expression levels calculated relative to vehicle control.

| Comparison | Fold Change | Confidence Intervals 65% | P-value | Result |
|---|---|---|---|---|
| SCX | | | | |
| Control (Vehicle) | 1 | 1 | | |
| Ad-GFP vs Ad-Smad3 | 1.879 | 1.347 - 2.598 | 0.059 | - |
| Ad-GFP vs Ad-GFP/T | 8.999 | 6.387 - 13.927 | 0.000 | UP |
| Ad-GFP vs Ad-Smad3/T | 12.000 | 7.304 - 18.210 | 0.002 | UP |
| Ad-GFP/T vs Ad-Smad3/T | 1.334 | 0.769 - 2.162 | 0.259 | - |
| FMOD | | | | |
| Control (Vehicle) | 1 | 1 | | |
| Ad-GFP vs Ad-Smad3 | 0.974 | 0.740 - 1.343 | 0.908 | - |
| Ad-GFP vs Ad-GFP/T | 3.531 | 2.428 - 5.268 | 0.000 | UP |
| Ad-GFP vs Ad-Smad3/T | 4.597 | 2.894 - 7.265 | 0.002 | UP |
| Ad-GFP/T vs Ad-Smad3/T | 1.302 | 0.827 - 1.890 | 0.202 | - |
| ADAMTSL2 | | | | |
| Control (Vehicle) | 1 | 1 | | |
| Ad-GFP vs Ad-Smad3 | 1.288 | 0.864 - 1.070 | 0.323 | - |
| Ad-GFP vs Ad-GFP/T | 23.292 | 16.119 - 35.690 | 0.018 | UP |
| Ad-GFP vs Ad-Smad3/T | 25.302 | 15.888 - 41.358 | 0.002 | UP |
| Ad-GFP/T vs Ad-Smad3/T | 1.086 | 0.561 - 2.105 | 0.602 | - |
| PRG4 | | | | |
| Control (Vehicle) | 1 | 1 | | |
| Ad-GFP vs Ad-Smad3 | 1.057 | 0.894 - 1.276 | 0.729 | - |
| Ad-GFP vs Ad-GFP/T | 3.620 | 3.079 - 4.336 | 0.002 | UP |
| Ad-GFP vs Ad-Smad3/T | 15.197 | 10.946 - 21.443 | 0.011 | UP |
| Ad-GFP/T vs Ad-Smad3/T | 4.199 | 3.146 - 5.498 | 0.022 | UP |

Table S7. qPCR details for Figure 4C. SCX, FMOD and ADAMTSL2 data was normalized to HPRT, expression levels calculated relative to TGFβ treated.

| Comparison | Fold Change | Confidence Intervals 65% | P-value | Result |
|---|---|---|---|---|
| SCX | | | | |
| TGFβ | 1 | 1 | | |
| T vs T/BIRB 1ug/ml | 1.208 | 0.227 - 6.467 | 0.586 | - |
| T vs T/ BIRB 5ug/ml | 1.221 | 0.212 - 7.246 | 0.568 | - |
| T vs T/ BIRB 10ug/ml | 1.362 | 0.214 - 9.20 | 0.552 | - |
| FMOD | | | | |
| TGFβ | 1 | 1 | | |
| T vs T/ BIRB 1ug/ml | 1.305 | 0.193 - 7.117 | 0.585 | - |
| T vs T/ BIRB 5ug/ml | 1.182 | 0.180 - 7.674 | 0.669 | - |
| T vs T/ BIRB 10ug/ml | 1.082 | 0.159 - 8.394 | 0.854 | - |
| ADAMTSL2 | | | | |
| TGFβ | 1 | 1 | | |
| T vs T/ BIRB 1ug/ml | 1.095 | 0.224 - 5.732 | 0.688 | - |
| T vs T/ BIRB 5ug/ml | 1.034 | 0.191 - 5.713 | 0.843 | - |
| T vs T/ BIRB 10ug/ml | 0.990 | 0.179 - 5.871 | 0.936 | - |

Table S8. qPCR details for Figure 4F. SCX, FMOD and ADAMTSL2 data was normalized to HPRT, expression levels calculated relative to TGFβ treated.

| Comparison | Fold Change | Confidence Intervals | P-value | Result |
|---|---|---|---|---|
| | | 65% | | |
| SCX | | | | |
| TGFβ | 1 | 1 | | |
| T vs T/MK 1ug/ml | 1.451 | 1.043 - 2.089 | 0.196 | - |
| T vs T/ MK 5ug/ml | 1.507 | 0.868 - 2.500 | 0.214 | - |
| FMOD | | | | |
| TGFβ | 1 | 1 | | |
| T vs T/ MK 1ug/ml | 1.258 | 1.048 - 1.580 | 0.123 | - |
| T vs T/ MK 5ug/ml | 1.338 | 1.110 - 1.514 | 0.016 | UP |
| ADAMTSL2 | | | | |
| TGFβ | 1 | 1 | | |
| T vs T/ MK 1ug/ml | 1.243 | 1.000 - 1.474 | 0.196 | - |
| T vs T/ MK 5ug/ml | 1.066 | 0.686 - 1.458 | 0.681 | - |

Table S9. qPCR details for Figure 5C. SCX, FMOD and ADAMTSL2 data was normalized to HPRT, expression levels calculated relative to TGFβ treated.

| Comparison | Fold Change | Confidence Intervals 65% | P-value | Result |
|---|---|---|---|---|
| SCX | | | | |
| TGFβ | 1 | 1 | | |
| T vs T/PD 1ug/ml | 0.821 | 0.192 - 3.300 | 0.684 | - |
| T vs T/PD 5ug/ml | 0.251 | 0.068 - 1.900 | 0.038 | DOWN |
| T vs T/PD 10ug/ml | 0.119 | 0.035 - 0.324 | 0.000 | DOWN |
| FMOD | | | | |
| TGFβ | 1 | 1 | | |
| T vs T/PD 1ug/ml | 0.871 | 0.191 - 3.985 | 0.769 | - |
| T vs T/PD 5ug/ml | 0.259 | 0.047 - 1.574 | 0.045 | DOWN |
| T vs T/PD 10ug/ml | 0.174 | 0.036 - 0.681 | 0.007 | DOWN |
| ADAMTSL2 | | | | |
| TGFβ | 1 | 1 | | |
| T vs T/PD 1ug/ml | 0.968 | 0.063 - 18.838 | 0.934 | - |
| T vs T/PD 5ug/ml | 0.266 | 0.049 - 1.450 | 0.034 | DOWN |
| T vs T/PD 10ug/ml | 0.119 | 0.021 - 1.205 | 0.006 | DOWN |

Table S10. qPCR details for Figure 5D. SCX data was normalized to HPRT, expression levels calculated relative to vehicle control.

| Comparison | Fold Change | Confidence Intervals | P-value | Result |
|---|---|---|---|---|
| | | 65% | | |
| EBF1 | | | | |
| Control (Vehicle) | 1 | 1 | | |
| Control (Vehicle) vs T | 0.589 | 0.418 - 0.834 | 0.000 | DOWN |
| T vs T/PD 1ug/ml | 1.067 | 0.983 - 1.159 | 0.518 | - |
| T vs T/PD 5ug/ml | 0.757 | 0.698 - 0.824 | 0.161 | - |
| T vs T/PD 10ug/ml | 0.994 | 0.519 - 1.916 | 0.663 | - |
| CMAF | | | | |
| Control (Vehicle) | 1 | 1 | | |
| Control (Vehicle) vs T | 0.622 | 0.556 - 0.696 | 0.000 | DOWN |
| T vs T/PD 1ug/ml | 0.863 | 0.717 - 1.052 | 0.323 | - |
| T vs T/PD 5ug/ml | 0.866 | 0.773 - 0.972 | 0.494 | - |
| T vs T/PD 10ug/ml | 0.842 | 0.657 - 1.092 | 0.331 | - |

Table S11. qPCR details for Figure 6B. SCX data was normalized to HPRT, expression levels calculated relative to Scrambled SiRNA control.

| Comparison | Fold Change | Confidence Intervals | P-value | Result |
|---|---|---|---|---|
| | | 65% | | |
| SCX | | | | |
| Scrambled siRNA | 1 | 1 | | |
| Scrambled siRNA vs Scx siRNA | 0.189 | 0.121 - 0.270 | 0.0001 | DOWN |

Table S12. qPCR details for Figure 6D. SCX, FMOD and ADAMTSL2 data was normalized to HPRT, expression levels calculated relative to Scrambled SiRNA control.

| Comparison | Fold Change | Confidence Intervals | P-value | Result |
|---|---|---|---|---|
| | | 65% | | |
| FMOD | | | | |
| Scrambled siRNA | 1 | 1 | | |
| Scrambled siRNA vs T/ Scrambled siRNA | 3.754 | 1.163 - 10.236 | 0.002 | UP |
| Scrambled siRNA vs Scx siRNA | 0.524 | 0.345 - 0.978 | 0.020 | DOWN |
| Scx siRNA vs T/ Scx siRNA | 2.155 | 0.815 - 5.054 | 0.074 | - |
| ADAMTSL2 | | | | |
| Scrambled siRNA | 1 | 1 | | |
| Scrambled siRNA vs T/ Scrambled siRNA | 3.462 | 1.012 - 11.618 | 0.029 | UP |
| Scrambled siRNA vs Scx siRNA | 0.998 | 0.369 - 3.397 | 0.999 | - |
| Scx siRNA vs T/ Scx siRNA | 2.148 | 0.480 - 7.983 | 0.203 | - |

Table S13. Primer sequences for qPCR in alphabetical order.

| Gene Name | Abbreviation | Forward Primer: 5'-3' | Reverse Primer: 5'-3' |
|---|---|---|---|
| Adamtsl2 | *Adamtsl2* | GGG-CAA-CAA-TCA-TCT-TGG-TTA-CT | CCG-TCG-GTA-CTT-GAC-CAC-T |
| C-Maf Proto-Oncogene | *cMaf* | GCT-TCA-GAA-CTG-GCA-ATG-AA | GTC-TCC-ACC-GGT-TCC-TTT-TT |
| EBF Transcription Factor 1 | *Ebf1* | GCA-TCC-AAC-GGA-GTG-GAA-G | GAT-TTC-CGC-AGG-TTA-GAA-GCC |
| Fibromodulin | *Fmod* | GGG-GTC-ACC-CAA-GCT-GCT-GT | TGA-CGT-CCA-CCA-CCG-TGC-AG |
| Hypoxanthine Phosphoribosyltransferase | *Hprt* | TCA-GTC-AAC-GGG-GGA-CAT-AAA | GGG-GCT-GTA-CTG-CTT-AAC-CAG |
| Proteoglycan 4 | *Prg4* | GAA-AAT-ACT-TCC-CGT-CTG-CTT-GT | ACT-CCA-TGT-AGT-GCT-GAC-AGT-TA |
| Scleraxis | *Scx* | ACT-CTT-CAG-TGG-CAT-CCA-CCT-TCA | TCT-GCC-TCA-GCA-ACC-AGA-GAA-AGT |

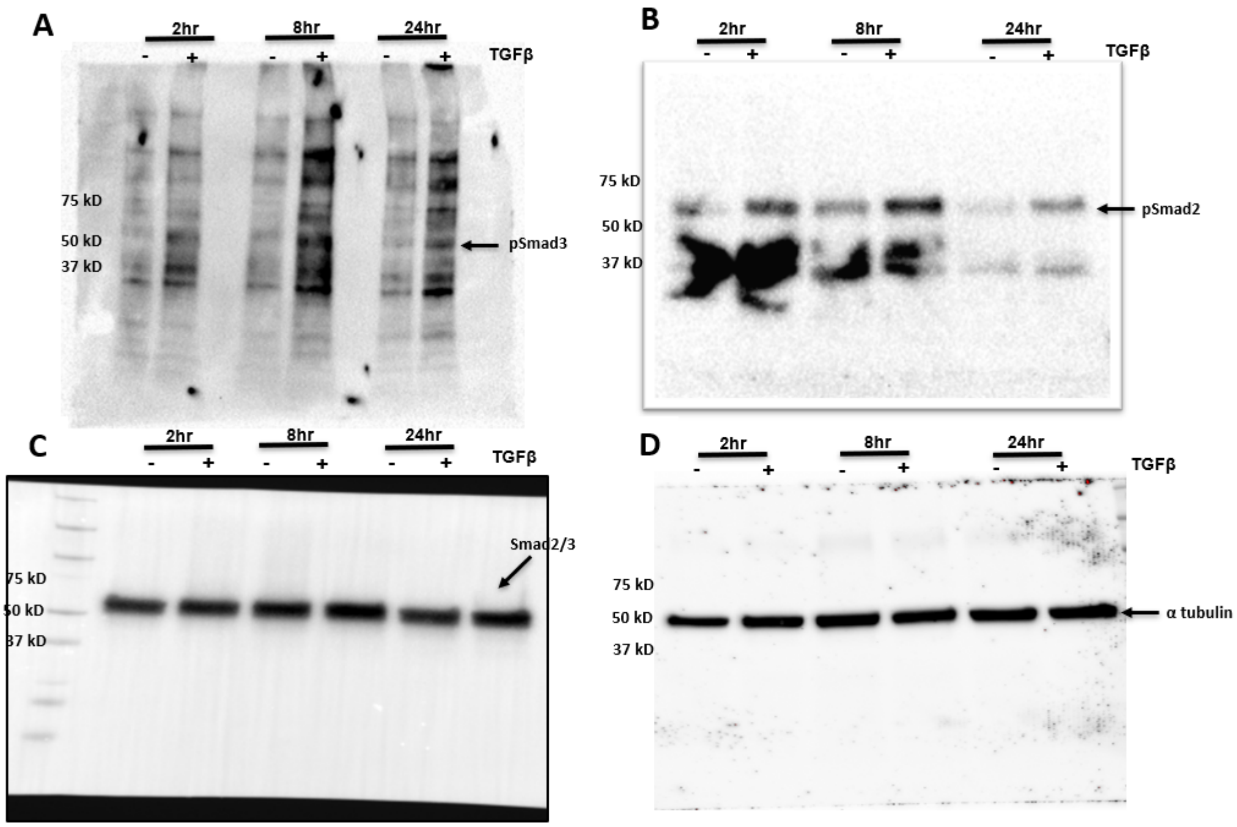

Figure S1. Example of uncropped (A) pSmad3 (B) pSmad2 (C) Smad2/3 and (D) α tubulin western blot from Figure 2, panel A

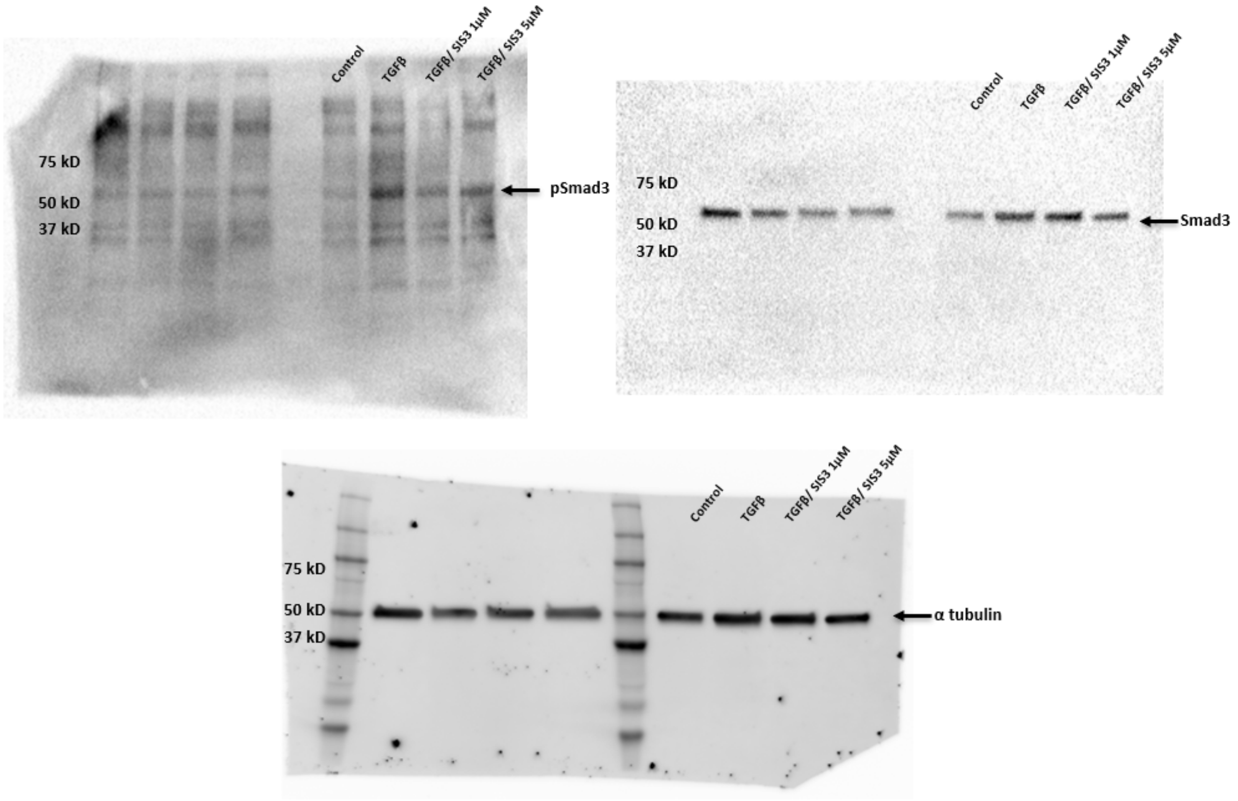

Figure S2. Example of uncropped (A) pSmad3 (B) Smad3 and (C) α tubulin western blot from Figure 2, panel D.

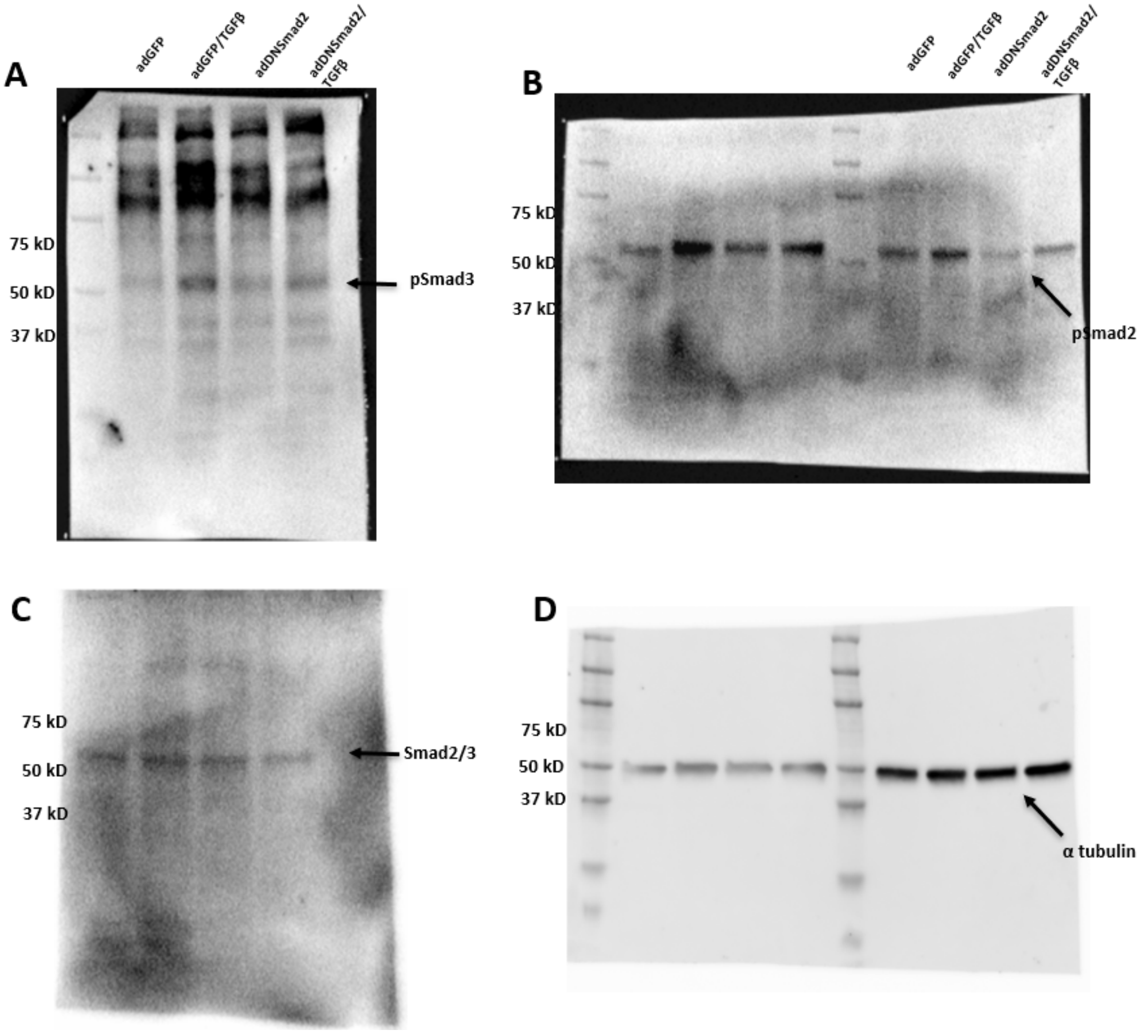

Figure S3. Example of uncropped (A) pSmad3 and (B) pSmad2 (C) Smad2/3 and (D) α tubulin western blot from Figure 2, panel G

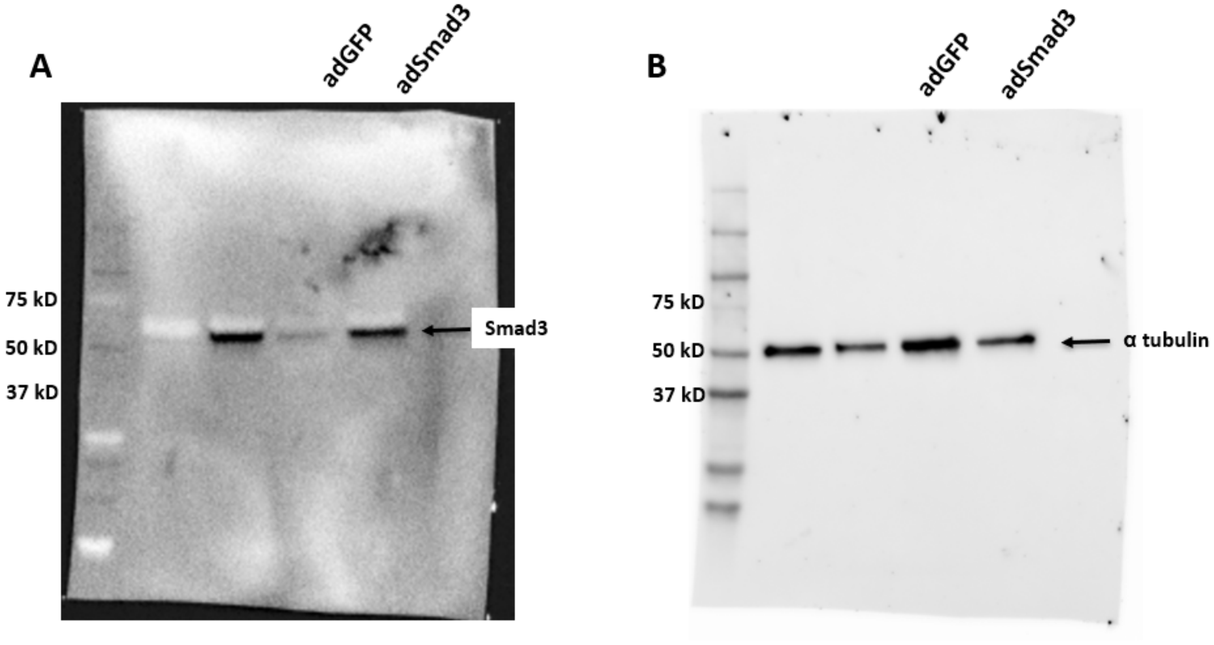

Figure S4. Example of uncropped (A) Smad3 and (B) α tubulin western blot from Figure 2, panel K.

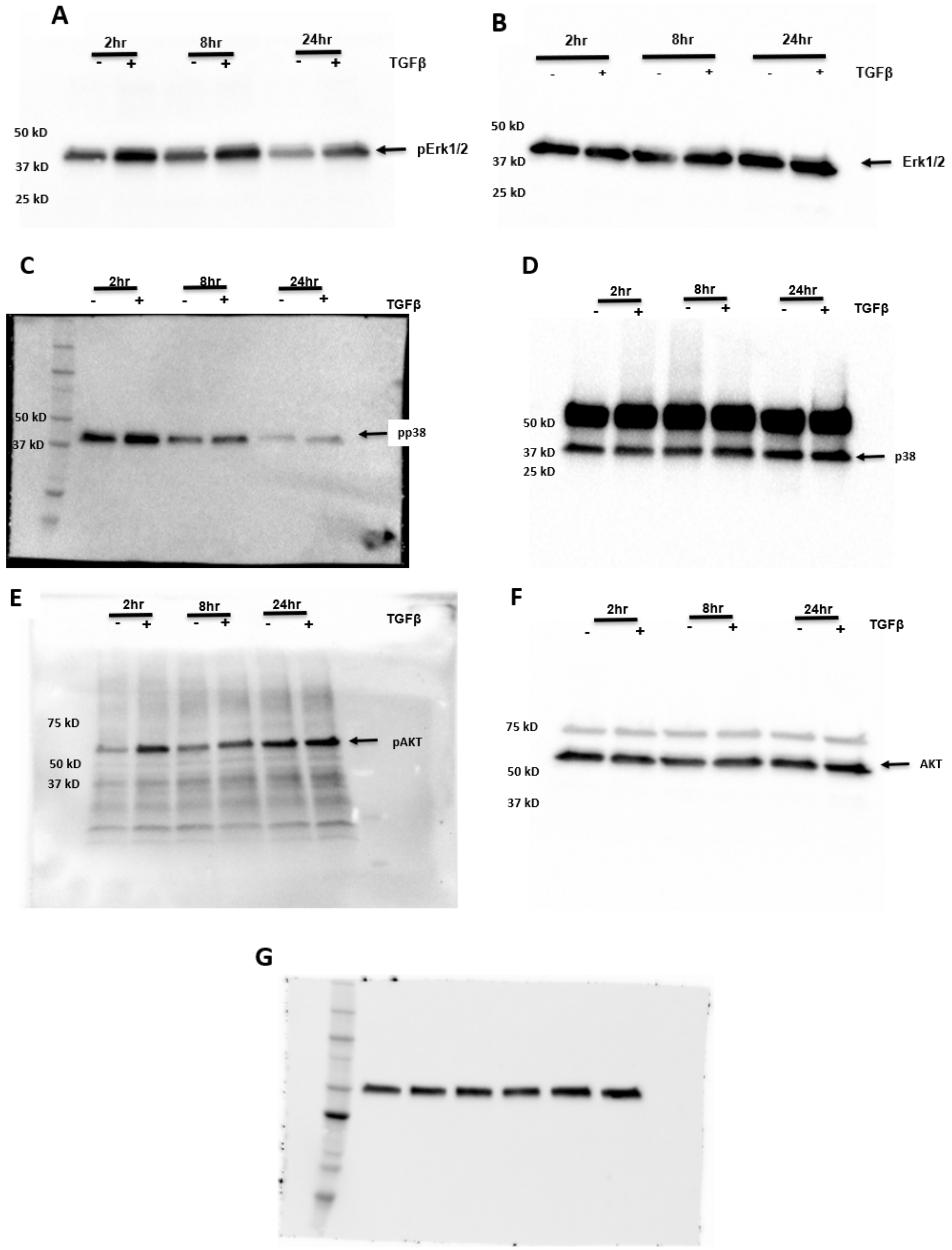

Figure S5. Example of uncropped (A) pErk1/2 (B) Erk (C) pP38 (D) P38 (E) pAKT (F) AKT and (G) α tubulin western blot from Figure 3, panel A.

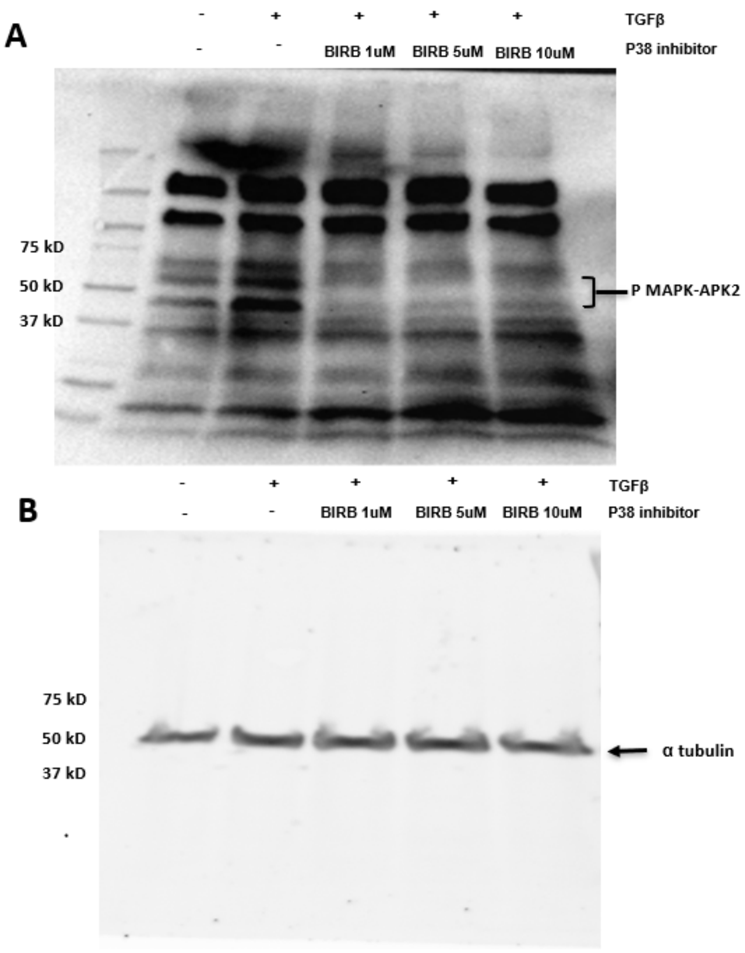

Figure S6. Example of uncropped (A) pMAPK-APK2 and (B) α tubulin western blot from Figure 4, panel A

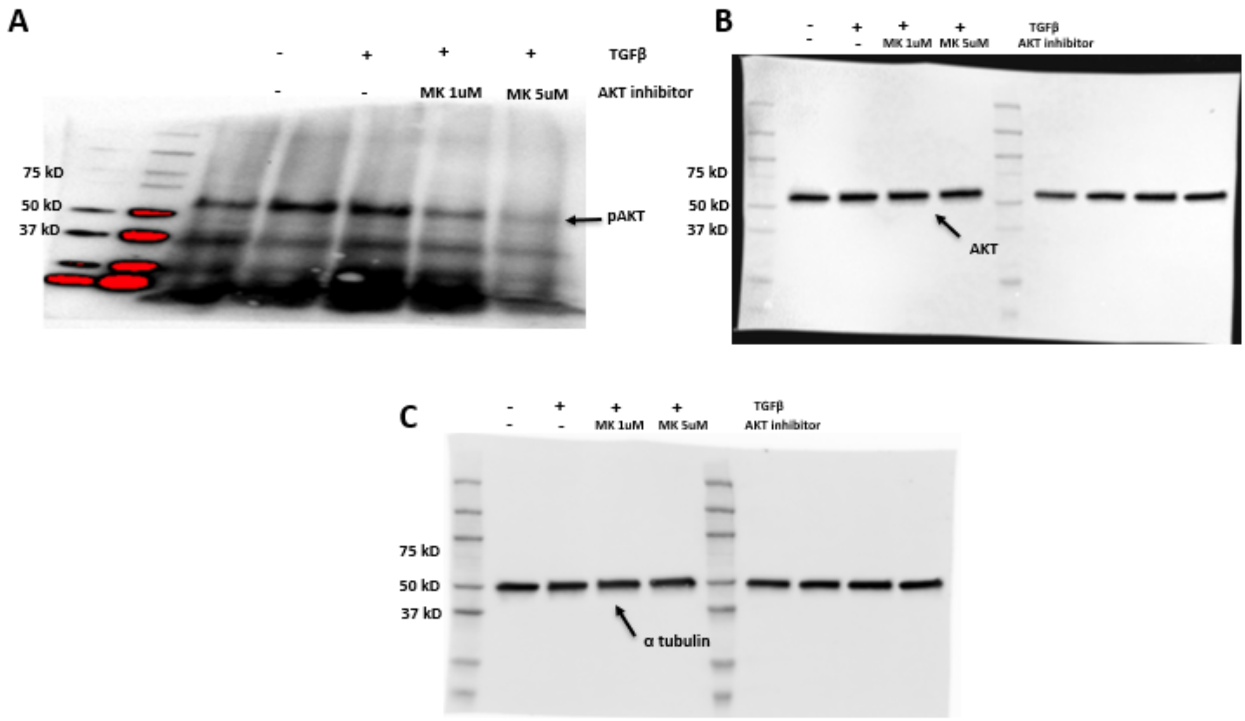

Figure S7: Example of uncropped (A) pAKT (B) AKT and (C) α tubulin western blot from Figure 4, panel D.

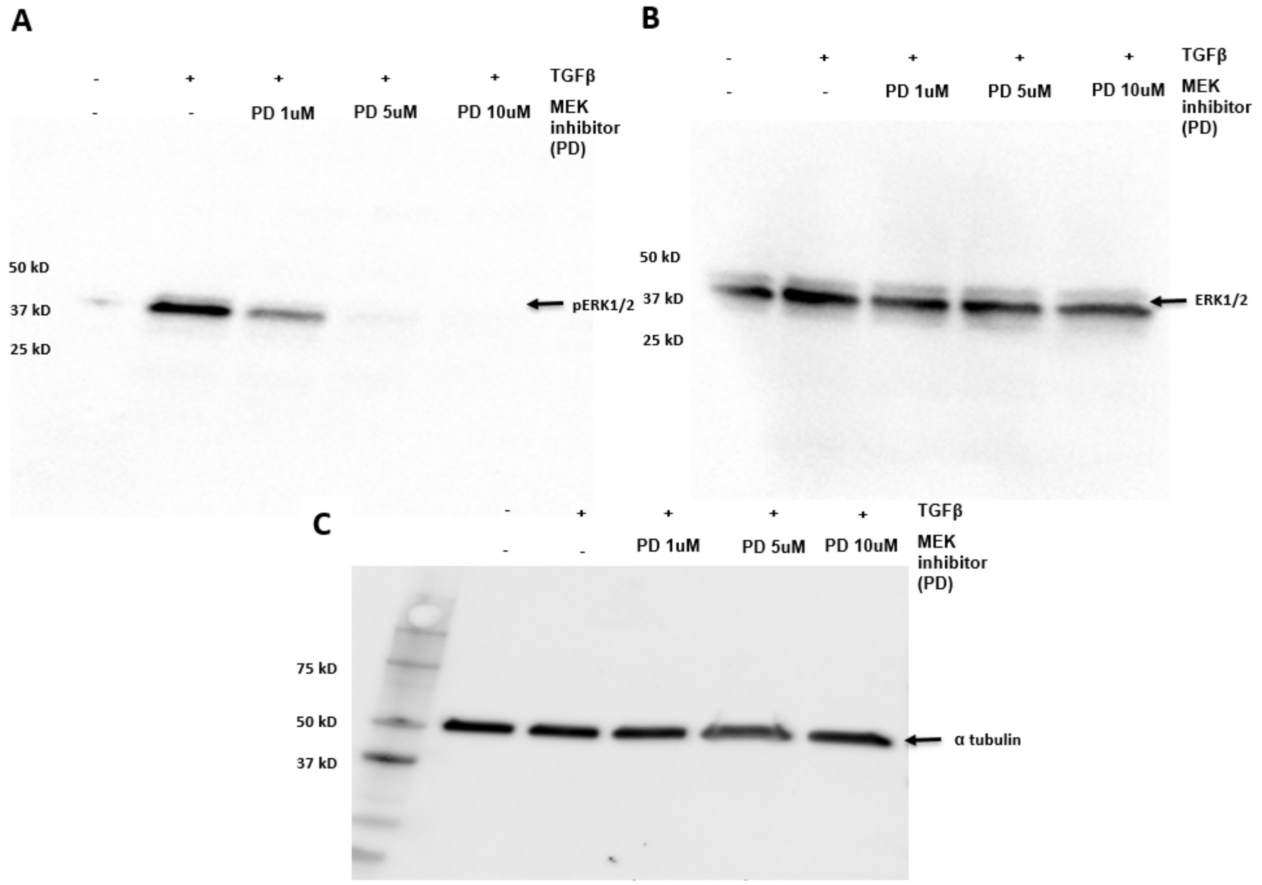

Figure S8: Example of uncropped (A) pERK1/2 (B) ERK1/2 and (C) α tubulin western blot from Figure 5, panel A.

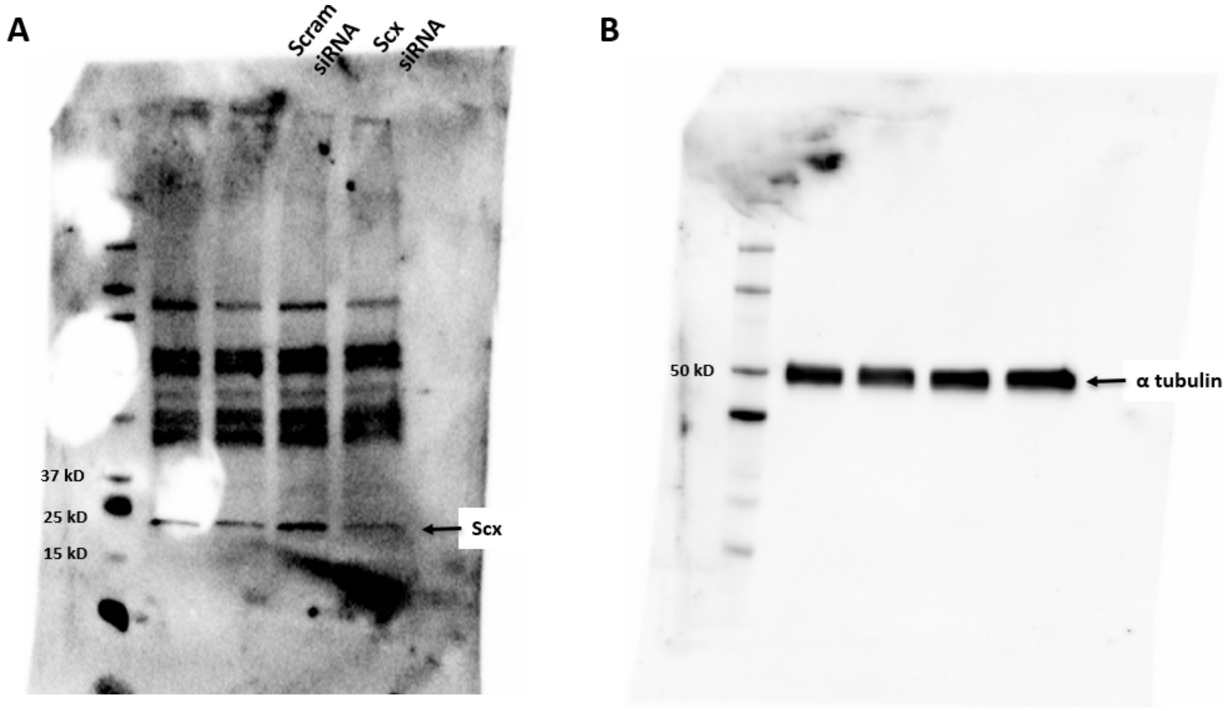

Figure S9: Example of uncropped (A) Scx and (B) α tubulin western blot from Figure 6, panel A.
